# XKidneyOnco: An Explainable Framework to Classify Renal Oncocytoma and Chromophobe Renal Cell Carcinoma with a Small Sample Size

**DOI:** 10.1101/2024.01.23.576782

**Authors:** Tahereh Javaheri, Samar Heidari, Xu Yang, Sandeep Yerra, Khaled Seidi, Mohammad Hadi Gharib, Tahereh Setayesh, Guanglan Zhang, Lou Chitkushev, Patricia Castro, Sayeeduddin Shahida Salar, Zahida Sayeeduddin, Neda Zarrin-Khameh, Mohammad Haeri, Reza Rawassizadeh

## Abstract

Renal oncocytoma and chromophobe renal cell carcinoma are two kidney cancer types that present a diagnostic challenge to pathologists and other clinicians due to their microscopic similarities. While RO is a benign renal neoplasm, ChRCC is considered malignant. Therefore, the differentiation between the two is crucial. In this study, we introduce an explainable framework to accurately differentiate ChRCC from RO, histologically. Our approach examined H&E-stained images of 656 ChRCC and 720 RO, and achieved a diagnostic accuracy of 88.2%, the sensitivity of 87%, and 100% specificity for explainable AI, which either outperforms or operate on par with convolutional neural network (CNN) models.

Besides, we enrolled 44 pathology experts (including pathologists and pathology trainees) to differentiate the two tumors. The average accuracy of pathologists was 73%, which is 15.2% lower than our framework.

These results indicate that the combination of human expert along with explainable AI achieve higher accuracy in differentiating the two tumors, while it reduces the workload of experts and offers the desired explainability for the medical experts.

## Introduction

Chromophobe Renal Cell Carcinoma (ChRCC) is the most common malignant renal neoplasm^(1)^. ChRCC subtype constitutes about 5-7% of all ChRCCs. A well-known benign mimicker of ChRCC, known as Renal Oncocytoma (RO), accounts for up to 6-9% of all adult renal tumors^(2)^. Proper differentiation between ChRCC and benign RO is very challenging task for pathologists, as they have similar histo-morphological features possibly because of their similar cell of origin known as intercalated cells of the collecting ducts ^(3)^.

A crucial step in this process is histologic examination. While the pathologic diagnosis can be straightforward for many cases of ChRCC with classic histology, the oncocytic variant of the ChRCC may look like an oncocytoma. These mimickers need additional studies for an accurate diagnosis. These studies have limited specificity, are costly, mostly limited to the tested tumor fragment, and highly time-consuming. New advances in fast and high-resolution slide scanning, along with machine learning techniques, have provided pathologists with unique opportunities to have a 24/7 expert assistant aid with a fast and accurate diagnosis.

Along with other medical fields, there is an increasing integration of computer-aided systems and automated tools in histopathology to enhance the precision of image interpretation. The emergence of explainable artificial intelligence (AI) in medical imaging holds promise to not only booster the accuracy and consistency of computer-aided diagnoses but also to amplify the trust and reliance placed by healthcare professionals on AI-driven medical solutions^(4-8)^.

Our study aims to explore four key research questions in this domain: (a) To what extent can an automated system assist pathologists in accurately diagnosing and distinguishing between RO and ChRCC, including hybrid cases? (b) Whether a Convolutional Neural Network (CNN) model, which is not explainable, may demonstrate superior accuracy compared to the explainable approach, while using small sample size, or (c) if a combination of CNN and explainable approach synergistically enhances our diagnostic accuracy? Finally, (d) What is the accuracy differences between human expert and our explainable framework?

To address these questions, we developed a high-precision, explainable, diagnostic framework based on traditional image preprocessing and human generated rules capabilities of OpenCV, a well-established image processing library^(9, 10)^. In addition, we implemented a CNN to juxtapose the outcomes of the explainable approach with deep neural networks. In our study, CNN models and explainable model were assessed on sets of H&E-stained images of 656 ChRCC and 720 RO. The explainable model demonstrated the following performance metrics: 87% sensitivity, 100% specificity, and 88.2% overall accuracy. Meanwhile, the best CNN model achieved 88% sensitivity, 90% specificity, and 88% accuracy. Finally, 21 images of ChRCC and RO were examined by 44 pathologists and pathology trainees, where they achieved an average accuracy of 73%. Sixteen percent of pathologist performed close or slightly better than AI at 81-90% accuracy, while around 48% of pathologist achieved an accuracy of 71-80%. The rest (38%) scored below 70% for the accuracy.

The performance metrics of the explainable approach operates on par to the CNN models, suggesting that a less computationally intensive explainable platform can match, and potentially exceed, the diagnostic accuracy of pathologists. Furthermore, our results demonstrate the capability to rival the performance of more complex systems such as CNN. The reason that we outperform the CNN originates in accurately identified rules designed by experts and this can manage the small cohort size of our study. In total, we have used only 1376 image, which is impractical to even fine-tune a pre-trained CNN model.

This strengthens the case for the explainable model as a reliable and robust assistance for diagnosing and differentiating renal tumors by pathologists. In particular, while leveraging a small dataset for training, our approach can assist pathologists in the challenging task of differentiating RO from ChRCC and hybrid tumors.

## Material & Methods

### Data

Images were acquired from the department of pathology at Baylor College of Medicine (Houston, TX) under institutional review board (IRB) approval H-40965.

In total, 656 ChRCC and 720 RO of H&E-stained images from 13 cases of oncocytoma and 15 cases of ChRCC were used to develop both explainable and deep-learning platforms.

## Results

The results section is organized into three primary subsections. First, we detailed the findings using explainable framework, followed by results from the CNN models. Lastly, outcomes from human assessments, conducted in parallel, to benchmark and compare with our approach is presented.

### Explainable Approach

In our effort to design and apply the explainable platform, the images ran through extensive histology image preprocessing steps. The preprocessing procedure comprises the following steps: (i) nucleus identification, (ii) cell membrane detection, (iii) noise removal and cell shape detection, (iv) cytoplasm intensity, (v) nucleus density and perinuclear halo space identification, (vi) automated report generation, and annotation.

### Histopathology Image pre-processing

Feature engineering is a crucial preprocessing step to develop an explainable approach or any other model that is not leveraging a neural network. However, due to known similarities between RO and ChRCC, identifying the most pertinent features posed real challenges. Table 1 explains distinct features of RO and ChRCC evaluated by a pathologist (i.e. MH). These features are generally accepted among pathologists and described in literature ^(11, 12)^.

**Table 1.**
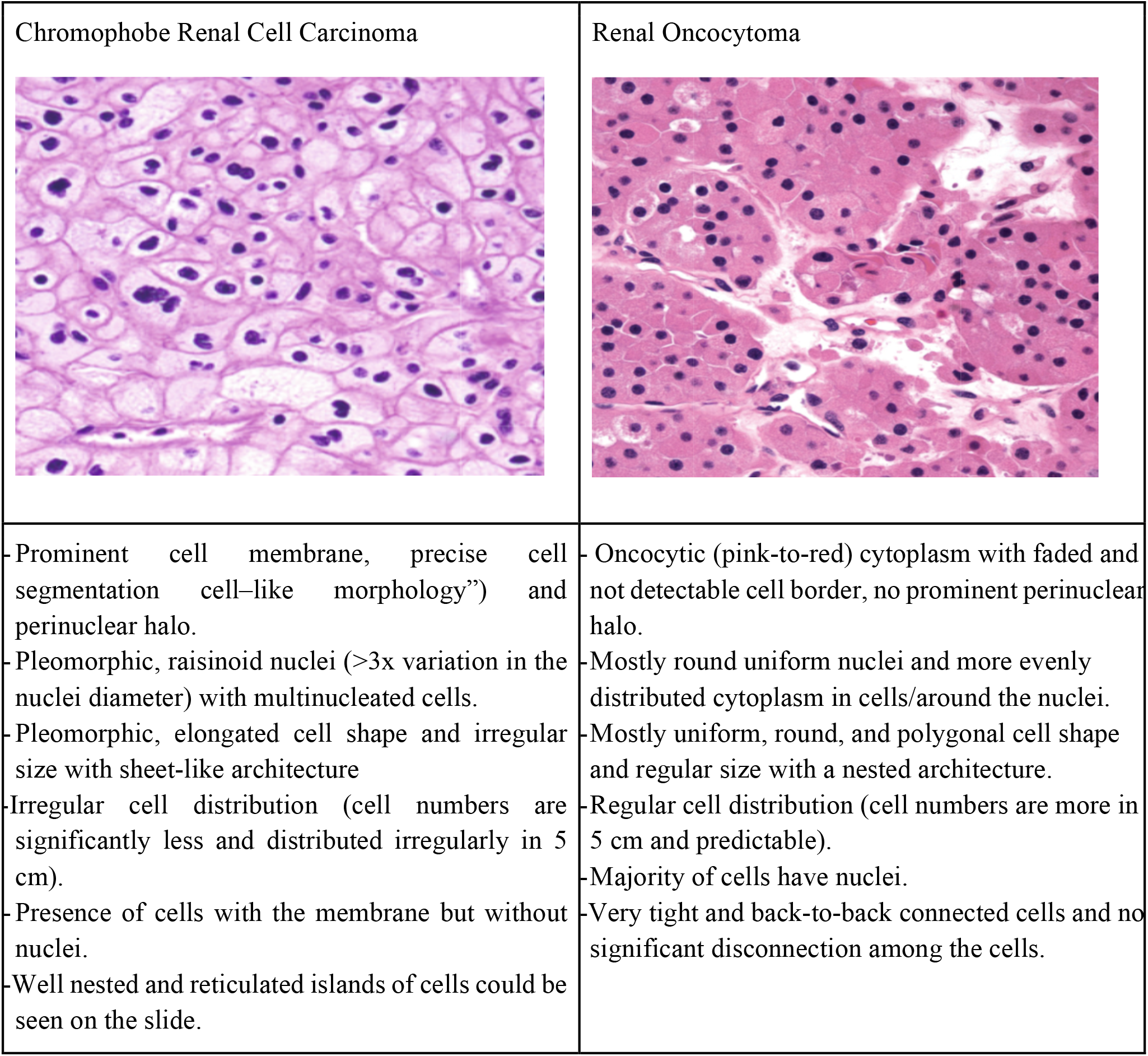
Presents the distinct features and differences of RO and ChRCC.

| Chromophobe Renal Cell Carcinoma | Renal Oncocytoma |
| --- | --- |
| - Prominent cell membrane, precise cell segmentation cell-like morphology”) and perinuclear halo.<br>- Pleomorphic, raisinoid nuclei (>3x variation in the nuclei diameter) with multinucleated cells.<br>- Pleomorphic, elongated cell shape and irregular size with sheet-like architecture<br>- Irregular cell distribution (cell numbers are significantly less and distributed irregularly in 5 cm).<br>- Presence of cells with the membrane but without nuclei.<br>- Well nested and reticulated islands of cells could be seen on the slide. | - Oncocytic (pink-to-red) cytoplasm with faded and not detectable cell border, no prominent perinuclear halo.<br>- Mostly round uniform nuclei and more evenly distributed cytoplasm in cells/around the nuclei.<br>- Mostly uniform, round, and polygonal cell shape and regular size with a nested architecture.<br>- Regular cell distribution (cell numbers are more in 5 cm and predictable).<br>- Majority of cells have nuclei.<br>- Very tight and back-to-back connected cells and no significant disconnection among the cells. |

Through extensive feature evaluation by expert pathologists, we determined the cell membrane, cell shape, cytoplasm color/intensity, and halo space to be the most discriminative and impactful features. These features consistently produced superior outcomes. To conduct our analysis, we employed the Gaussian Blur^(13)^, Gray Scaling, Binarization and Erosion features^(14)^ implemented by OpenCV image processing library^(9, 10)^, as depicted in Figure 1.

**Figure 1.**
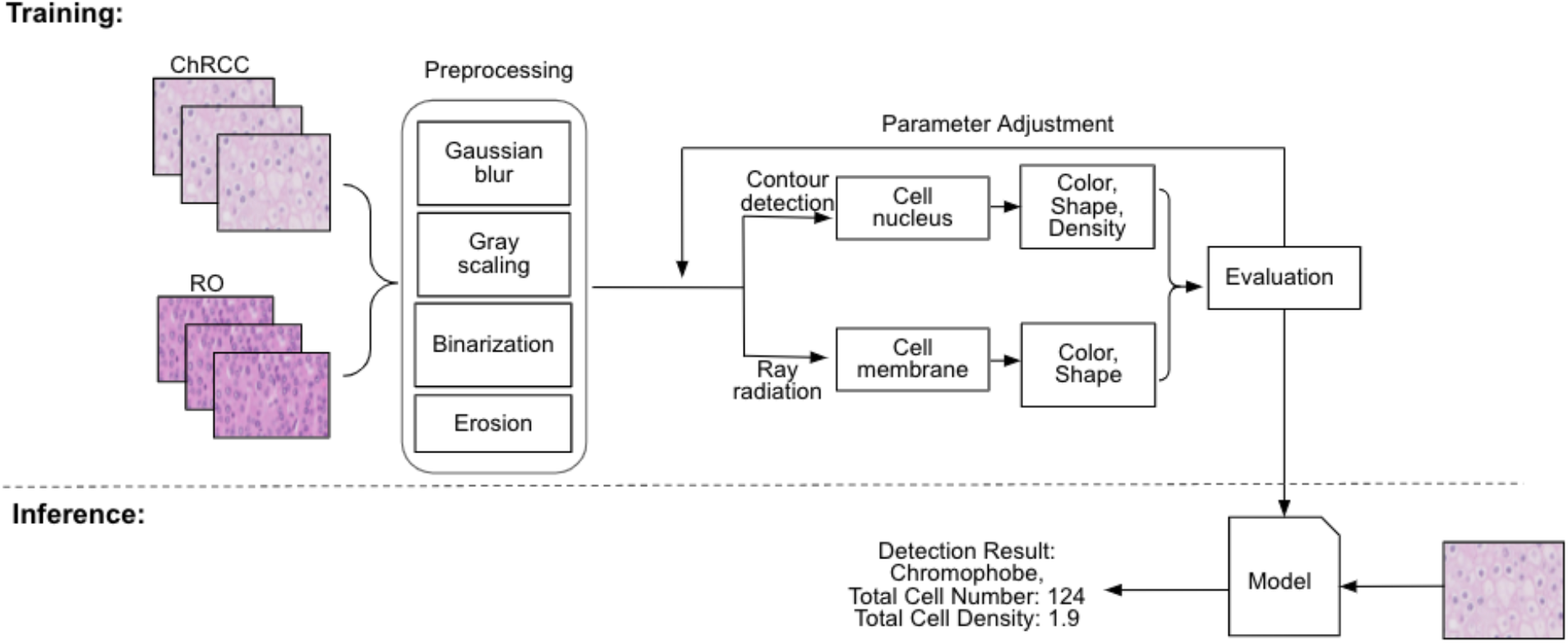
The process flow of proposed framework for RO and ChRCC image classification.

As it is shown in Figure 1, the preprocessing steps are used to build train the model. Later while the model is ready, we use the preprocessing steps as well. However, for the sake of understanding, we did to present them in the inference stage.

#### 1. Nucleus identification

The nuclei identification was one of the crucial features of the explainable approach, as underscored by differences between ChRCC and RO nuclei illustrated in Figure 2a. To pinpoint the nucleus, we applied a trio of standard image preprocessing steps: grayscale conversion, gaussian blur, and image erosion. The subsequent step involved setting a binarization threshold to underscore the nucleus while filtering out unnecessary elements like the cell membrane. This thresholding ensured that any pixel values between [0, 255] were either designated as 0 (black, indicating the nucleus) or 255 (white, indicating the surrounding area of the nucleus). A sharp increment in black pixel proportions, represented by an elbow point in each curve (Fig 2b), highlights the most significant transition point in pixel ratios.

**Figure 2.**
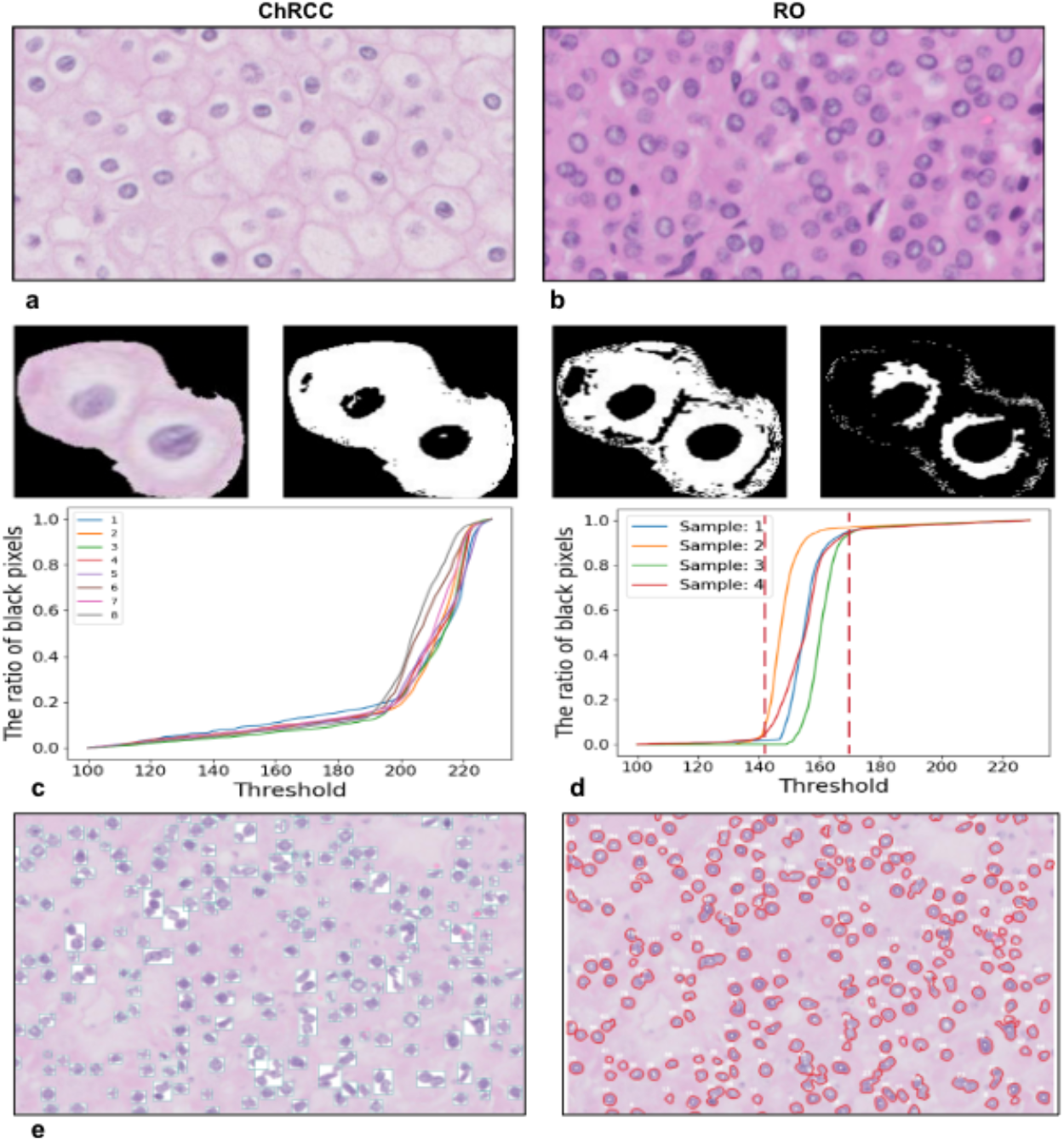
Highlighting the Significance of Nucleus Intensity in Image Analysis. **(a)** Original images showcasing ChRCC (left) and RO (right). **(b)** Depicts two ChRCC cells, illustrating the process of threshold setting and display the outcomes of thresholding at values of 185, 200, and 215, respectively. **(c-d)** Illustrating the rising proportion of black pixels as the threshold value escalates. The x-axis represents the threshold value, while the y-axis indicates the ratio of black pixels. **(e)** Image after binarization (left); the processed image with marked areas (right).

Given the varied Hematoxylin and Eosin (H&E) stain color variations in oncocytoma and chromophore images, we abandoned a rigid preprocessing protocol. Instead, we iteratively tested threshold values, ranging from 155 to 230, in increments of 5. This method aimed to identify the threshold by rendering the maximum cell count. For instance, a suboptimal threshold would either under-represent the nucleus or erode the cell’s intrinsic shape and roundedness, leading to its wrong exclusion or inclusion. Our threshold parameter sensitivity analysis, which spanned thresholds from 160 to 205 (Fig 2d), was corroborated by cell detection. It ensures that pixels exceeding the set threshold are identified as black (indicating cell), with the rest labeled white.

Characteristic cellular distinctions are evident when contrasting RO (Fig 2c) and ChRCC cells (Fig 2d). Specifically, the delayed rise in ChRCC cell values signifies their characteristically lighter cytoplasmic hue. Following threshold establishment, we refine the native image into a rendition seen in Fig. 2e, left. Each nucleus is subsequently assigned a unique identifier (ID), prepping them for subsequent border delineation.

#### 2. Cell membrane detection

The distinctiveness of the cell membrane is a primary feature in differentiating ChRCC from RO^(11, 15)^. To accurately detect the cell membrane, we devised an algorithm that casts rays outwardly from the center of a nucleus towards the adjacent cell nuclei. The algorithm initiates by projecting a central vertical ray (illustrated as the thicker line in Fig3a) and subsequently casts additional rays at intervals of 5 degrees, totaling 72 rays. For illustrative simplicity, Fig3a displays only 12 of these rays and not all of them.

As each ray traverses through the grayscale image, pixel values are computed. Given the pathway of the ray through the nucleus, cytoplasm, and cell membrane, varying pixel values are encountered. When the distance from the nucleus center exceeds 24 (denoted as point B in Fig 3b), there’s a notable increase in color depth. This lowest color depth (point B) is identified as the cell membrane location, indicating the point where a ray intersects the neighboring cell. The trigonometric function was used to determine the endpoints of the ray and the ray coordinates.

**Figure 3.**
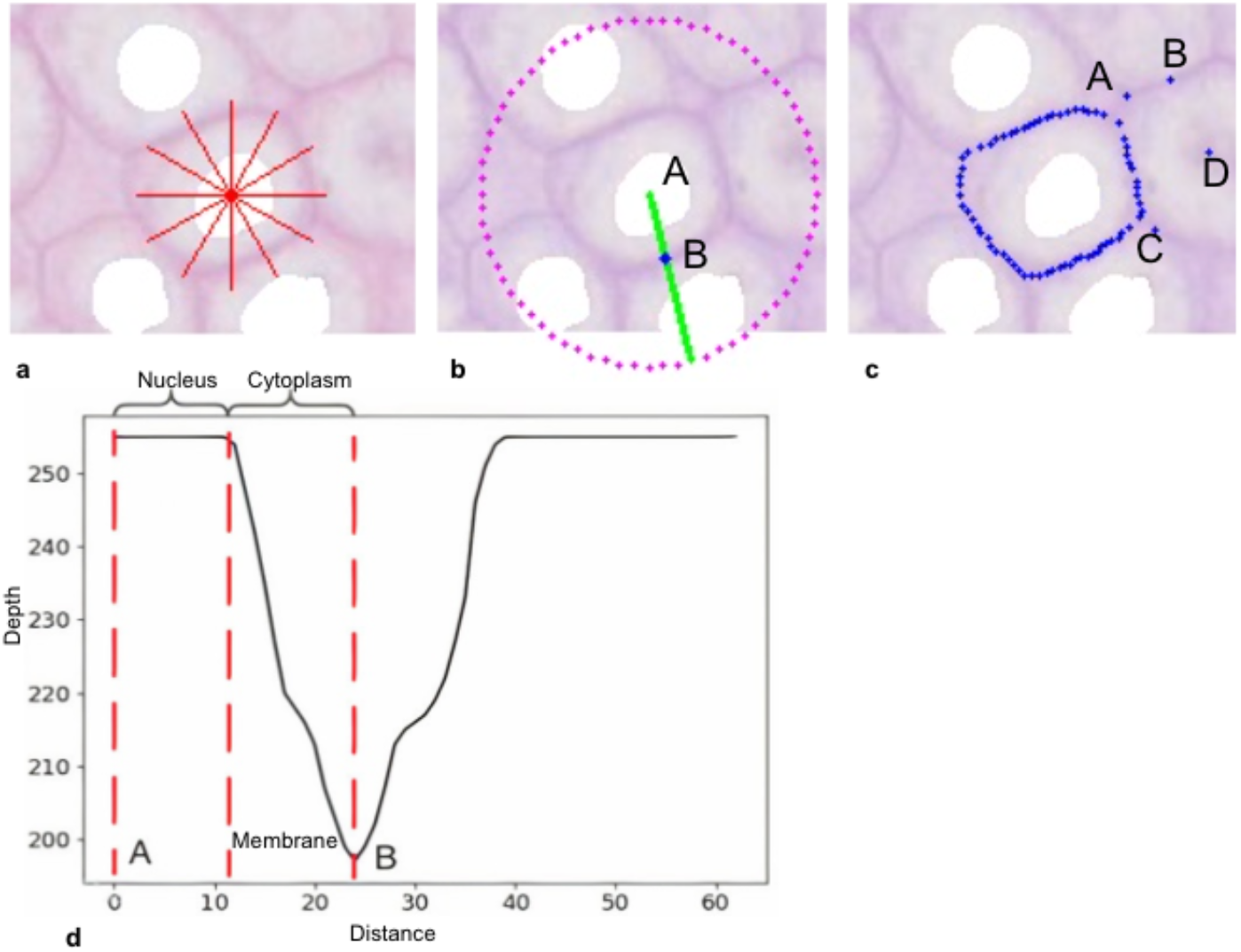
Ray-based Cell Membrane Detection. **(a)** Representation of 12 equidistant rays, each 30° apart, originating from the cell center (Note: 5° spacing used in actual computations). **(b)** Point b is detected and generated by algorithm as the crises point when the ray faces the membrane. **(c)** Highlighted regions A-D indicate algorithmic false positives in cell membrane identification. **(d)** Profiling pixel intensity (grayscale) as a function of distance from the cell center in a plot.

Upon analyzing the entire set of 72 rays, as visualized in Figure 3c, the deepest color depth for each ray was calculated. In Fig 3c, these membrane intersections are highlighted using blue dots in the image. As shown in Fig 3d, certain data points (labeled A, B, C, and D) appeared anomalous, suggesting noise, which we address in the subsequent noise-removal phase.

#### 3. Noise (anomaly) removal and cell shape detection

Our approach employs a common noise reduction, signal smoothing via averaging. In particular, it averages the values of adjacent data points when a spike in distance to the center is detected. Given the inherent continuity of cell boundaries, anomalies are indicative of noise (Fig 3d). The interrupted points (A-D) and anomalies in continuity are shown in Fig. 3d, and they are presented in Fig. 4a with respect to their distances from identified points and the cell nucleus. Points exhibiting substantial deviation from their neighboring points are counted as noise. Fig. 4d shows the refined cell membrane in both ChRCC and RO. A continuous cell membrane is characteristic of ChRCC (Fig. 4d, Left), whereas a fragmented cell membrane aligns with RO’s features (Fig. 4d, Right). The distinct morphological differences, with ChRCC displaying an elongated, spindle-shaped cellular form and RO a rounded, polygonal morphology, further enhance the precision of the differentiation between the two. The red dot line is the original data, and the yellow dot line is the average of points, and the line is supposed to serve as a standard circle. The green line is the cell membrane generated by our algorithm after adding the standard circle. To calculate the difference between each cell’s distance and the standard circle’s radius, the algorithm uses the following equation: 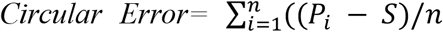, In this equation, *n* represents the number of rays, *P*_*i*_ represents the distance between each point and the center of the circle, and *S* represents the standard radius, which is the average distance of all the points from the center of the circle. The formula calculates the difference between each point’s distance and the standard radius and then sums up these differences divided by the number of rays.

**Figure 4.**
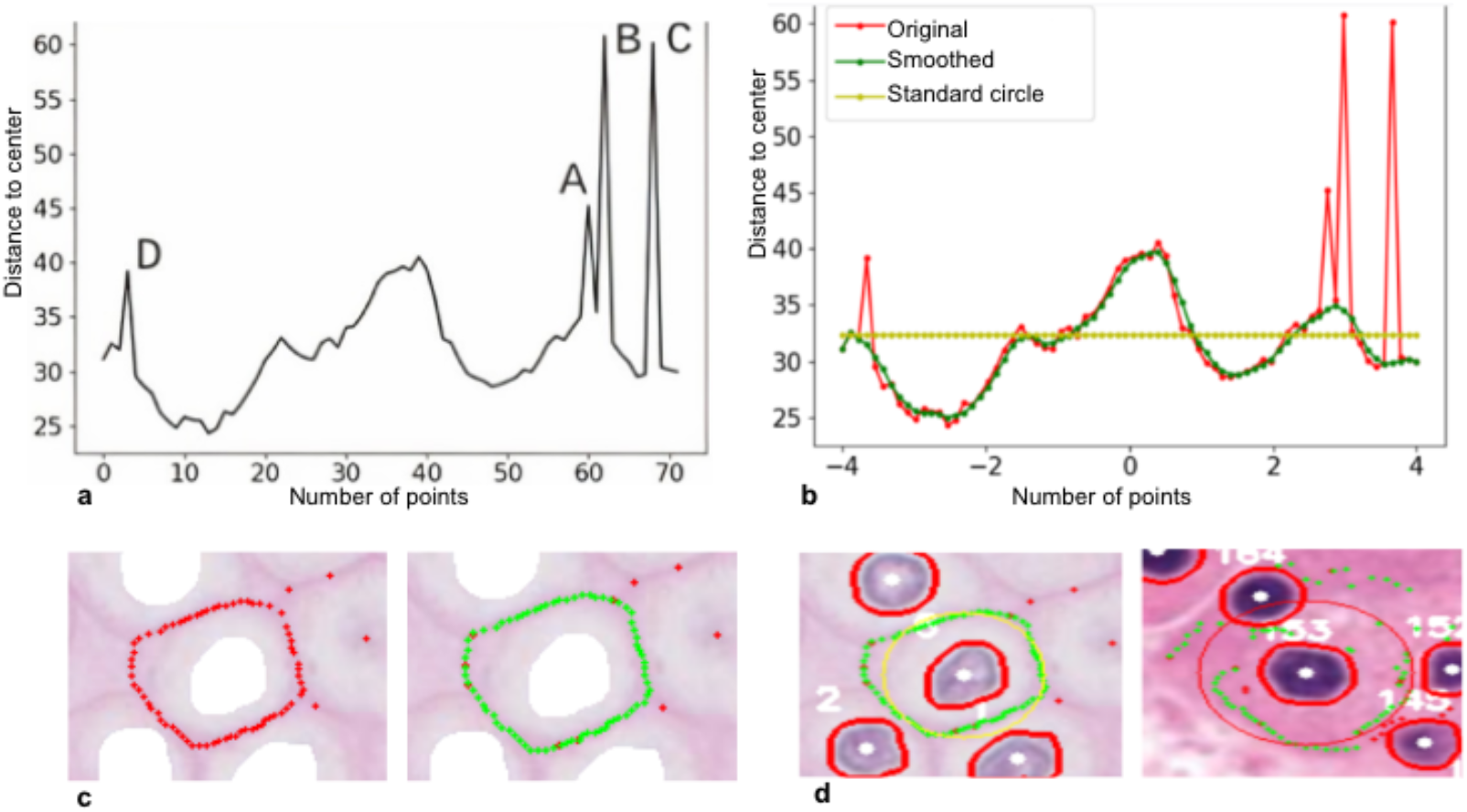
Noise detection and removal in cell membrane identification. **(a)** Illustrates interruptions in the continuity of cell boundaries, signaling potential noise or anomalies. **(b)**. graphical representation of the distances between highlighted points (red in 4c) and the cell nucleus. Points A-D, mark distances from their neighbors, are identified as noise.**(c)**.The cell depiction with identified points in red, used for assessing the noise based on their radial distances from the cell center. **(d)** Comparison of refined cell membrane identification in ChRCC (left) and RO (right).

#### 4. Cytoplasm intensity

To delineate color intensity variations, our algorithm traces a path between the centers of adjacent nuclei. The ChRCC (Fig. 5a left) is characterized by its paler cytoplasm and nuclei, resulting in an RGB signature (Fig. 5b) distinct from that of the RO cell (Fig. 5c). The notable decline in the ChRCC RGB curve (Fig. 5c) signifies a pronounced cell membrane. Fluctuations in color density, as depicted between Fig. 5c-d, manifest as oscillations in the RGB curve, termed changing points. Our refined algorithm is tailored to detect and capture these abrupt minima and maxima in the curve.

**Figure 5.**
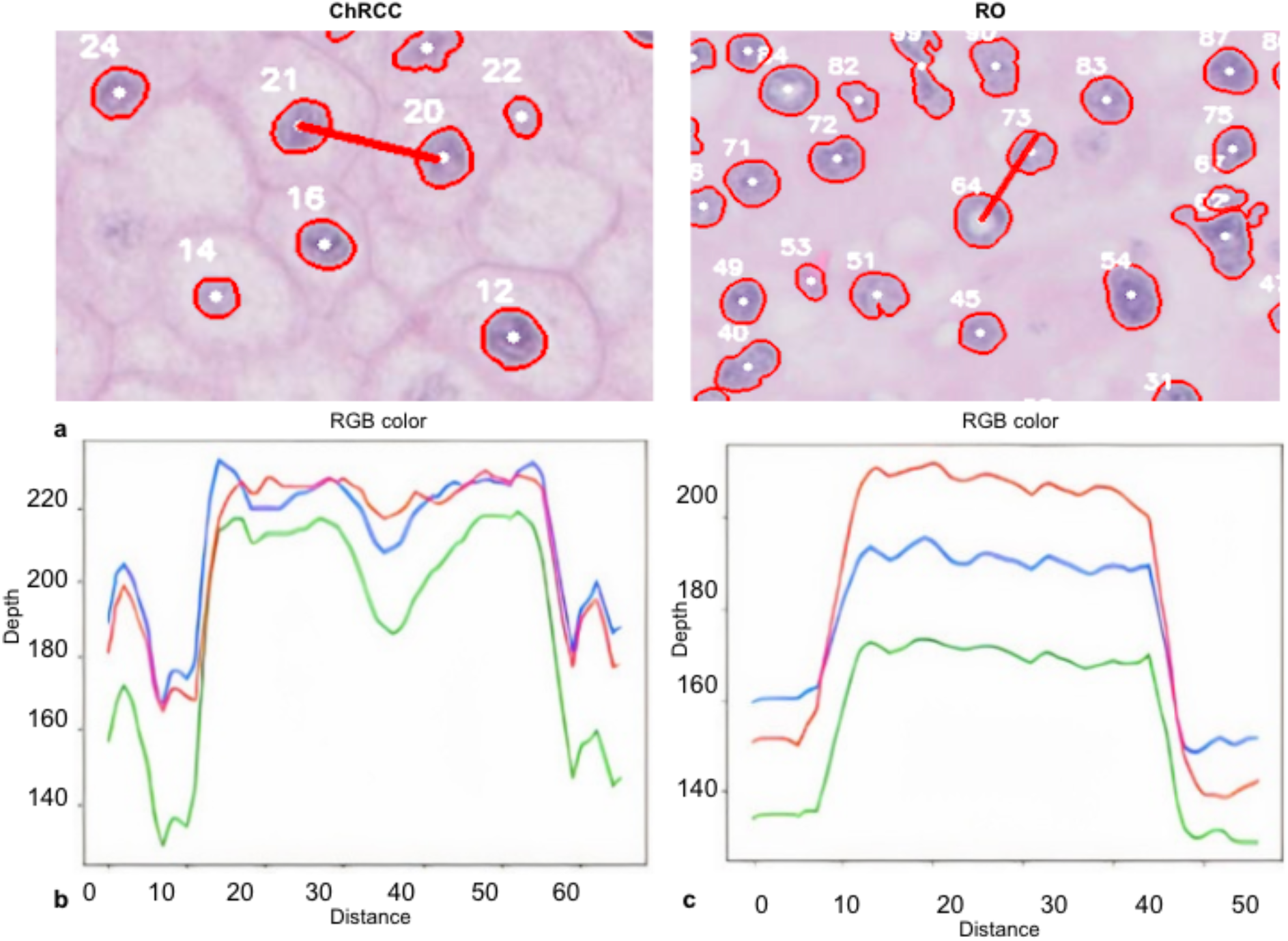
Cytoplasm Intensity Analysis in Chromophobe and RO Cells. **(a)** Microscopic visualization of the chromophobe (left) showing its characteristic lighter cytoplasm and nuclei compared to RO (right). **(b)** The RGB curve for the chromophobe, with a marked decline, indicates a distinct cell membrane. **(c)** Variations in the RGB curve represent changing points caused by shifts in color density.

**Figure 6.**
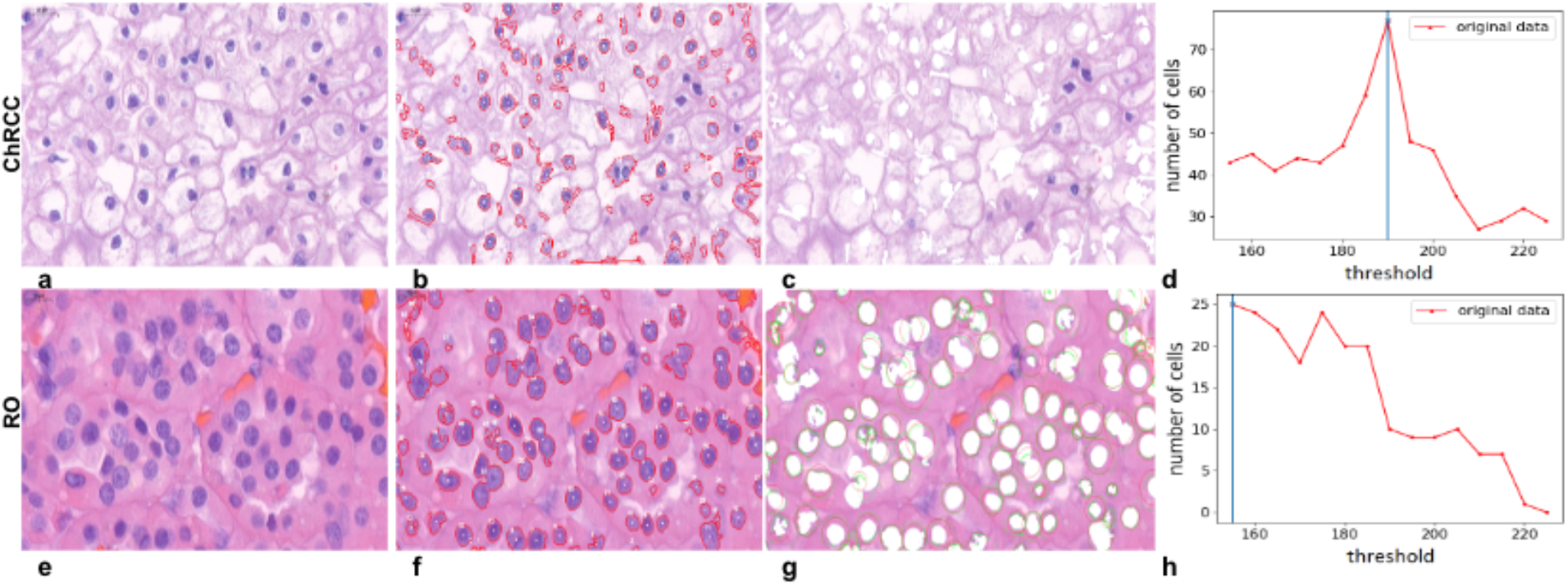
Perinuclear halo analysis in ChRCC and RO cells. **(a)** and **(e)** display the ChRCC and RO cells. **(b), (c), (f)** and **(g)** represent binarization threshold test highlighting the disparate color distributions in ChRCC compared to RO cells. **(d)** and **(h)**, ChRCC profile requires a higher initial threshold due to its distinct perinuclear halo and uneven cytoplasmic content.

#### 5. Nucleus density and perinuclear halo space identification

Nuclear morphology is still an important indicator that distinguishes the two types of cancers. The perinuclear halo, a distinct white space surrounding the nuclei, is a characteristic feature of ChRCC cells. This feature leads to noticeable color distribution variations between ChRCC and RO cells. In ChRCC, the nucleus is surrounded by clear halo spaces, whereas in RO the nucleus is solid and darker (fig6 a and e, ChRCC and RO, respectively). After running experimental comparisons (during our formative evaluation), we found that there was a difference in the mean color depth between two types of cells. Upon applying a binarization threshold test (fig6 b, c, f and g), the initial turning point in the ChRCC profile necessitates a higher threshold than that of the RO cells. As the threshold value increases, an expansive shift in color depth becomes evident, underscoring the irregular cytoplasmic content, including the perinuclear halo, in ChRCC cells.

#### 6. Automated report generation and annotation

All the steps of the above-mentioned framework bring us to a simplified and streamlined functionality of our framework: the generation of an automated report. As depicted in Figure 7, when a pathological image is processed, the framework produces an annotated image with a concise textual summary of the results. This ensures that the insights derived from our analysis are explainable and easy to interpret by expert, facilitating their application in diagnostic settings.

**Figure 7.**
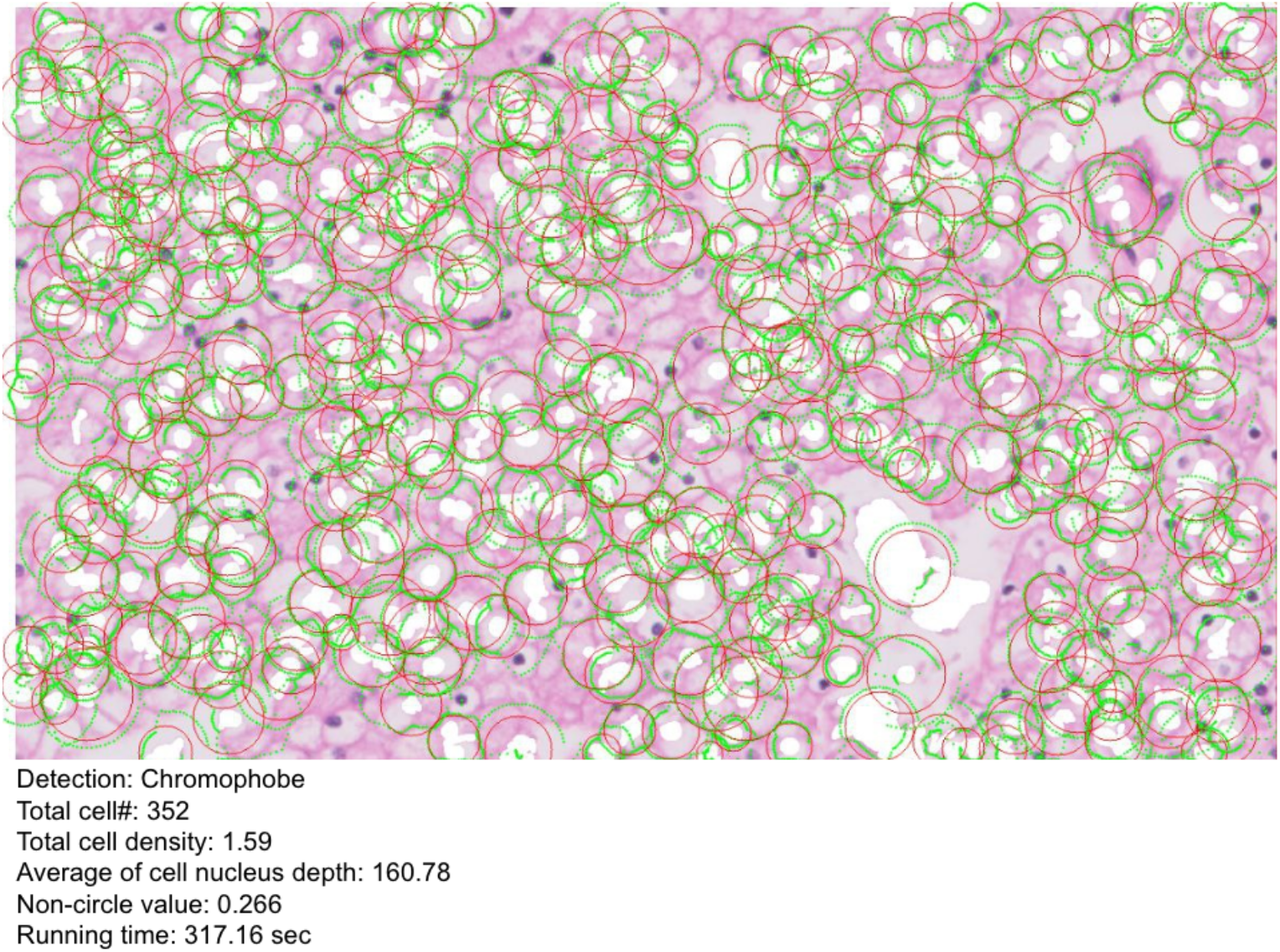
Automated single cell report generation that consolidates the results from our analysis. Upon processing a pathological image through our framework, an annotated image is automatically produced.

### Neural Network Approach

Our study incorporated different convolutional neural network models to juxtapose the performance of an explainable AI methodology against a fully autonomous classification framework^(16)^. To intensify the specificity of our analysis, we employed a patching approach, dissecting tumors into numerous sub-regions for close-up scrutiny. This micro-focused strategy is particularly advantageous for identifying distinct types of cancer cells, shown in Figure 8.

**Figure 8.**
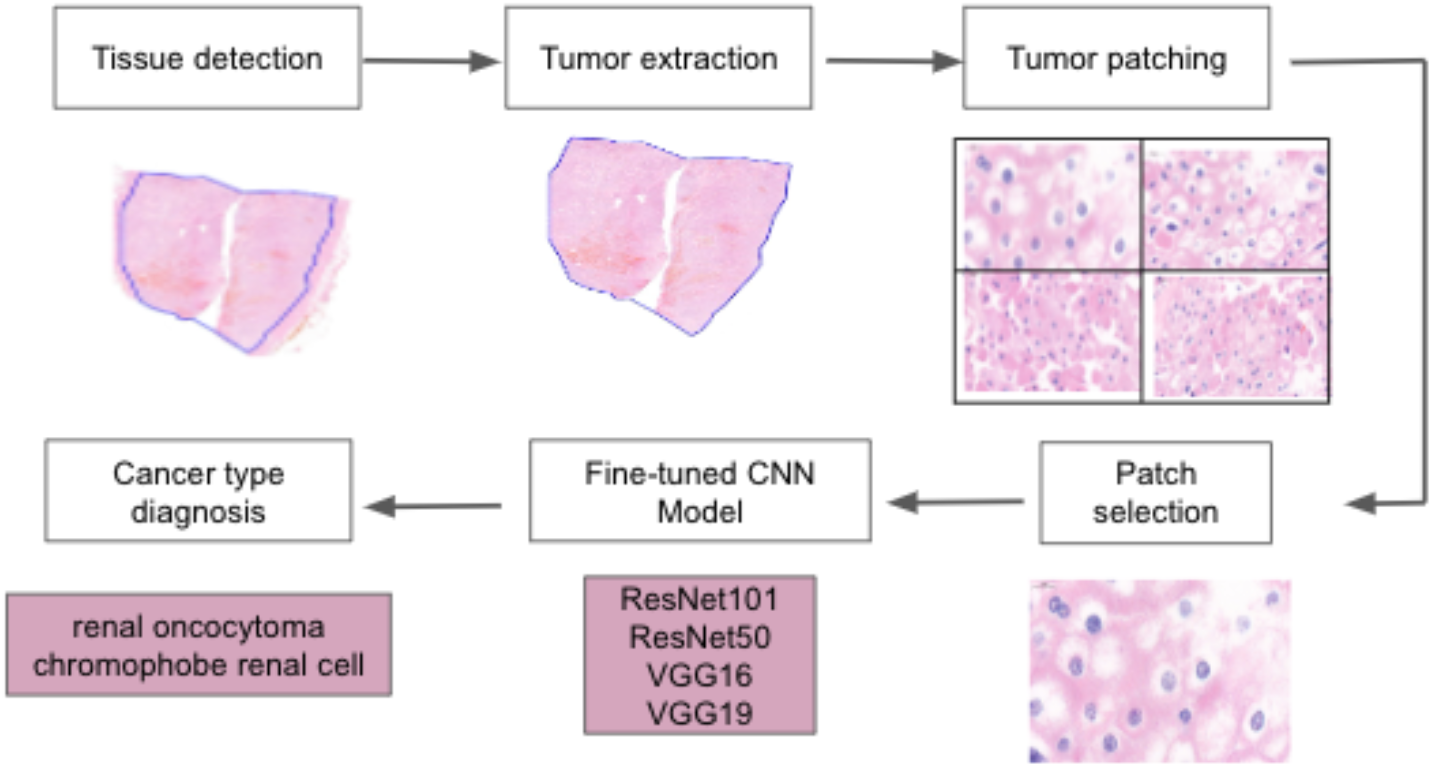
Workflow of analysis with neural network. This figure presents a schematic representation of the patching and classification process used to differentiate between chromophobe and oncocytoma cells in kidney tumor samples.

It’s imperative to note that the neural network models were fine-tuned to differentiate RO and ChRCC cells, deliberately excluding non-cancerous components such as vascular structures, adipose cells, and intravascular blood cells.

The patching process, while useful, is not without its challenges. Certain patches might inadvertently encompass peripheral areas of the tumor, which may harbor less cancer cells. These boundary regions produce patches with a mix of cancer and non-cancer elements, muddling the model’s output and diminishing predictive accuracy.

First, we used UNET and MaskRCNN for patching, which resulted in low accuracy as 86.3% and 85.1% for each, respectively^(17, 18)^. To counteract this, we adopted a hands-on approach to patch selection in these ambiguous zones, improving the model’s precision. The dimension of the patches is also a factor of significant consequence. Optimal patch sizes are essential for a nuanced understanding of cellular morphology and mitotic activity. However, the inherent size variability within ChRCC and RO cells complicates the standardization of patch dimensions. Despite the availability of automated object detection methods such as the Feature Pyramid Network^(19-21)^, our strategy favored human experts for patch selection.

Our pathologist meticulously identified 40 patches from each chromophobe specimen and selected approximately 51 images from each oncocytoma sample, to ensure a rich cellular detail is given to the model. After the patches were selected carefully, they were fed into three different pre-trained CNN classifiers ResNet101, ResNet50, VGG19 and VGG16. As a result, those models are fine-tuned with our image samples.

Figure 8 illustrates the procedural workflow of the using the fine-tuned CNN models and table 2 enumerates the results yielded by three distinct classifiers.

**Table 2.** Comparative performance of classifiers in tumor analysis. This table summarizes the results of three different classifiers used to discriminate between chromophobe and oncocytoma cells within kidney tumor samples.

| Model/Classifier | Accuracy | Specificity | Sensitivity |
| --- | --- | --- | --- |
| VGG16 | 82% | 84% | 82% |
| VGG19 | **88%** | 90% | **88%** |
| Resnet50 | **88%** | 90% | **88%** |
| ResNet100 | 82% | 88% | 81% |
| Explainable | **88%** | **100%** | 87% |

Compiling our data, the dataset consists of 1440 images - an even split between chromophobe and oncocytoma. Data distribution was slated at 70% for training, 20% for validation, and the final 10% for test.

In the development of our model for ChRCC, we utilized 466 images for training, 121 for validation, and 69 for testing. For Renal RO, the corresponding distribution was 492 images for training, 153 for validation, and 75 for testing.

Table 2 consolidates the performance metrics (accuracy, specificity, and sensitivity) of the explainable AI and the four additional classifiers deployed in the deep learning platform. These metrics reflect the outcomes from the final testing phase, which was conducted on the images evaluated by the pathologists. The overall accuracy achieved among pathologists in this study was 73%.

### Human assessment

Forty-four pathologists and pathology trainees were asked to differentiate ChRCC from renal oncocytoma (RO) by examination of images prepared at a magnification comparable to 200X with an inset image comparable to 400X.

Twenty-one images were selected by a pathologist (M.H.) from 7 cases of ChRCC and 8 cases of renal oncocytoma, initially generated from H&E-stained slides. Image selection was based on presence of the tumor in more than 70% of the image, minimal out-of-focus regions present in the image, absence of any artifacts, proper and uniform H&E stain and absence of folded tissues. Nine images of RO and 12 images of ChRCC were included in the survey with the imbalance to prevent bias. The images were limited to 21 for practical considerations to encourage careful examination of the image without fatigue. Our preliminary studies suggested that more than 25-30 images can lead to loss of interest to make the accurate differentiation. There was no time limit to finish the survey.

The participants were recruited by email invitations to do the survey. All participants were instructed to look at the tutorial images of the two tumors at the beginning of the survey to become familiar with the morphology and staining of each tumor. Following the initial step, they were asked to review each image and vote for either the ChRCC or RO. The total number of ChRCC and RO cases was not released to participants to prevent bias. The input from each participant was recorded for further analysis.

## Discussion

Pathologic diagnosis in histopathology is the process of reaching a diagnosis by microscopic examination of cells and tissues from the sampled specimen. Pathologic diagnosis is a complex process starting with reviewing the patient’s medical history, imaging and laboratory studies, among other diagnostic report before the procedure of tissue sampling. The examination of the tissue during the sampling procedure is usually performed to reassure proper tissue sampling. Pathologists examine the hematoxylin and eosin stain (H&E) slides and would request additional sections, additional immunohistochemistry and/or special stains, and/or additional molecular studies based on standard guidelines. The final diagnostic description will be issued after assembling all the data into the final integrated pathology report.

Despite its similar histomorphology to RO, the ChRCC is considered a malignant renal tumor while RO is benign. This poses a challenge for pathologist to accurately differentiate the two and report it to clinicians. New molecular signatures are described in RO, ChRCC and also in hybrid oncocytic/chromophobe tumors (HOCT). Accordingly, the chromosomal alteration in RO can be null or with 1, 14, 21, Y deletions. The ChRCC demonstrates, null or a wide range of chromosomal alterations (1, 2, 3, 5, 6, 9, 10, 11, 13, 17, 21, Y del Other mutations and varied RNA expression profiles have also been reported^(22)^. However, there is no reliable molecular method for accurate differentiation of the two tumors.

Explainable artificial intelligence are computer algorithms and models trained to provide answer along with a human understandable explanation. On the other hand, computer assisted diagnosis have shown promising results in other cancer types such as breast and lung cancer diagnoses^(23, 24)^. We utilized similar approach to assist differentiation of malignant ChRCC and benign RO; which have different clinical management. We reviewed several cases with diagnoses of ChRCC and RO in our institution between 2001 and 2016. Multiple images from different foci of each tumor were captured and analyzed by image processing software performing nuclear segmentation followed by detecting nearest-neighbor, nuclear shape/size and nuclear densities/area algorithms. Overall score for each was calculated, analyzed and compared^(25)^. In that study, nuclear segmentation step approached ∼94% accuracy for both ChRCC and RO using binary mode or Fourier transform/band pass filter setting. Cell boundaries detection showed similar results. A scoring system utilizing a combination of nearest-neighbor, nuclear shape/size and nuclear densities/area was also used and showed ∼93% accuracy in differentiation between ChRCC and RO in well-fixed/prepared section. While this approach held promise for good results, however the diagnostic accuracy was reduced when the image contained normal renal tissue adjacent to the tumor. This was an approach to utilize traditional machine learning for detection of histological features associated with specific type of tumors to reach an accurate diagnosis based on accepted criteria.

This evidence-based computer aided diagnosis explains the steps of reaching to a diagnosis. In this study, a complete different explainable AI was employed in parallel with a conventional Convolutional Neural Network (CNN) to enhance the precision of tumor differentiation. CNN was used as it already confirmed the liability in challenging cases and studies^(26)^. Operating concurrently, these two models yield diagnostic results independently, without reliance on external systems. This combination increases the precision while still has the histological evidence for making the diagnosis.

We further compared the pathologists and the framework for differentiation of the two tumors and showed that our framework outperformed human experts in making an accurate diagnosis. However, a group of pathologists managed to become as good as machine in terms of precision, when they collaboratively analyze the image. Computer assisted diagnosis can be used as an ancillary tool to differentiate ChRCC from renal RO and to reduce the cost of immunohistochemical stains. Our current image processing algorithm has managed to differentiate ChRCC and RO with high accuracy in well-fixed/prepared sections. Adoption of additional nuclear/cellular features to modify this algorithm will improve the specificity of this method.

To our knowledge, this is the first work of utilizing an image processing algorithm to differentiate between the two ChRCC and RO kidney cancers by an explainable AI showing the histological features leading to the diagnosis.

Our framework stands out not just for its accuracy but also for its reasonable reporting pace. It highlights the exact location and boundaries of each nucleus and differentiates between RO and ChRCC. Unlike other approaches, this allows pathologists to directly verify specific areas, streamlining their review process.

## Conclusion

In conclusion, our study XKidneyOnco: An Explainable Approach to classify Renal Oncocytoma and Chromophobe Renal Cell Carcinoma with a small sample size has demonstrated the potential of an explainable AI to assist pathologists in the challenging task of differentiating between RO and ChRCC. Despite the histological similarities that complicate the diagnosis, our explainable framework, supported by a CNN, achieved high diagnostic accuracy, sensitivity, and specificity. The rule based model alongside with CNN model outperformed individual pathologist assessments and presented a reliable alternative to more resource-intensive molecular methods. Furthemore, our study success in applying leveraging to a relatively small dataset underscores the models’ robustness and the viability of this method in clinical practice. Despite high accuracy of neural network models, they require large dataset for training.

Our approach maintains transparency in its decision-making process, which is crucial for clinical acceptance. The insights gained from this research pave the way for further development of AI-assisted diagnostic tools in nephropathology. Such tools can provide valuable second opinions, reduce the time and cost associated with traditional human-only diagnostic methods, and ultimately, enhance patient care by ensuring accurate and timely diagnoses. Our findings advocate for the continued integration of computer aided diagnosis in pathology.

